# Adipose fin development and its relation to the evolutionary origins of median fins

**DOI:** 10.1101/283432

**Authors:** Thomas A. Stewart, Robert K. Ho, Melina E. Hale

## Abstract

The dorsal, anal and caudal fins of vertebrates are proposed to have originated by the partitioning and transformation of the continuous median fin fold that is plesiomorphic to chordates. Evaluating this hypothesis has been challenging, because it is unclear how the median fin fold relates to the adult median fins of vertebrates. To understand how new median fins originate, here we study the development and diversity of adipose fins. Phylogenetic mapping shows that in all lineages except Characoidei (Characiformes) adipose fins develop from a domain of the larval median fin fold. To inform how the larva’s median fin fold contributes to the adipose fin, we study *Corydoras aeneus* (Siluriformes). As the fin fold reduces around the prospective site of the adipose fin, a fin spine develops in the fold, growing both proximally and distally, and sensory innervation, which appears to originate from the recurrent ramus of the facial nerve and from dorsal rami of the spinal cord, develops in the adipose fin membrane. Collectively, these data show how a plesiomorphic median fin fold can serve as scaffolding for the evolution and development of novel, individuated median fins, consistent with the median fin fold hypothesis.

## Introduction

Fins have evolved repeatedly in vertebrates^1-­4^ and, thus, provide a powerful system for studying how new body parts originate. Primitively, chordates are characterized by a median fin fold (MFF), a midline structure comprised of dorsal and ventral portions that meet posteriorly to form a protocercal tail^2^. The extinct chordates *Haikuichthys* and *Haikuella* exhibit this condition, with the ventral portion of the LMFF interrupted by the anus^5, 6^. The extant cephalchordate amphioxus also has a MFF, which passes to the right of the anus uninterrupted^7^. Spatially differentiated, individuated median fins evolved later, in craniates^1^. These new fins are hypothesized to have originated by the partitioning of the MFF into multiple fin modules^4, 8-­12^. Specifically, the dorsal, anal and caudal fins are predicted to have evolved from the MFF by its reduction in some positions and its retention in others. This ‘median fin-­fold hypothesis’ is related to the ‘lateral fin-­fold hypothesis’ of paired pectoral and pelvic fin origin, which itself posits that paired continuous fins along the flank were subdivided to create the pectoral and pelvic fins^8-­10^. Although the lateral fin-­fold hypothesis has largely been abandoned in favor of a scenario where pectoral fins evolved first and pelvic fins evolved secondarily^1, 13, 14^, the MFF hypothesis remains influential.

In many fishes, ontogeny appears to recapitulate the phylogenetic transformational scenario predicted by the MFF hypothesis. For example in zebrafish, *Danio rerio* (Cyprinidae), a larval median fin fold (LMFF) encompasses the trunk early in development^15^. The LMFF develops as the somites are forming;; specification and outgrowth proceeds in a caudal-­to-­rostral direction, driven by *Fgf* signaling^15^. The LMFF is composed of an epithelial bilayer medial to which are actinotrichia (tapered collagenous rods organized approximately parallel to the fin’s proximodistal axis), which sandwich a core of mesenchyme^16^. Later in development, spatially discontinuous adult median fins—the dorsal, anal, and caudal fins—form^17, 18^, and the LMFF is reduced by apoptosis in positions that do not bear adult fins^19^.

Adult median fins are described as developing from the LMFF^15, 20-­24^. However, *D. rerio* mutants suggest that the development of adult median fins does not depend on proper formation of the LMFF^25^. Further, most tissues that comprise adult fins (*e.g.*, dermal and endoskeleton, musculature, and fin-­associated innervation) are not derived from tissues in the LMFF, but from other sources, including paraxial mesoderm^20, 26, 27^. Thus, while the LMFF might function as scaffolding for the morphogenesis of adult fins (*e.g.*, actinotrichia guiding the migration of osteogenic mesenchyme that forms lepidotrichia^28, 29^), the relationship between the LMFF and adult fins is not one of straightforward ontogenetic transformation. This poses a challenge to recapitulist^30^ arguments for the MFF hypothesis^8-­10^.

Here, to inform hypotheses of (1) phylogenetic transformation from MFFs into individuated fins and (2) ontogenetic transformation of the LMFF into adult fins, we study the diversity and development of adipose fins. These appendages have evolved repeatedly within teleosts^3^ and are positioned on the dorsal midline between the dorsal and caudal fins. Adipose fins have been studied as models of how form and function evolves in vertebrate appendages^3, 31-­^ ^33^ and might also inform how development evolves to generate novel appendages. Descriptions of adipose fin morphogenesis are scattered throughout the literature—in taxonomies of larval fishes, staging papers for select taxa, and a study of early development of these fins^34^. We aggregate the data on adipose fin development from the literature and analyze them in a phylogenetic context. Additionally, we characterize adipose fin development in the South American armored catfish *Corydoras aeneus* (Gill 1858) (Siluriformes, Callicithyidae), focusing on the development of the adipose fin skeleton and sensory anatomy. Collectively, these data reveal that adipose fins can evolve and develop by retention and elaboration of a domain of the LMFF. We discuss how these data inform hypotheses of median fin origin in early vertebrates.

## Results

### Diversity of adipose fin development

Analysis of the literature yielded information on adipose fin development for twenty-­four species belonging to five orders of fishes (Suppl. Table 1). Two patterns of adipose fin development are observed, consistent with previous descriptions^34, 35^ (Fig. 1 a). In Characoidei (Characiniformes), adipose fins develop *de novo* as buds following the reduction of the LMFF. In two Characoidei genera, *Brycinus* and *Phenacogrammus,* the adipose fin grows out before the LMFF has been completely reduced^34^. In all other clades for which data is available, adipose fins appear to develop by the retention of a domain of the LMFF between the dorsal and caudal fin (Fig 1 b).

**Figure 1.**
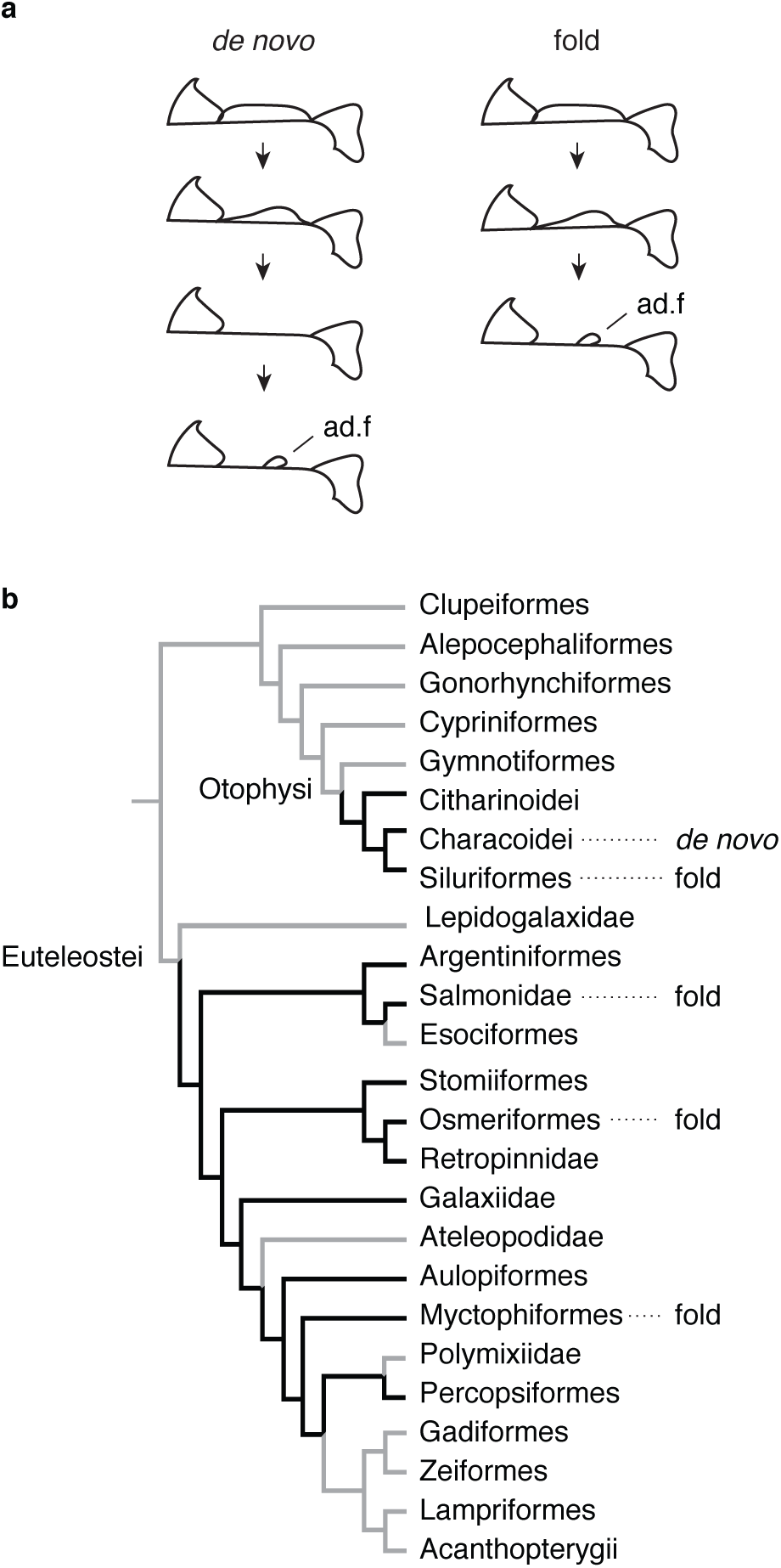
Distribution of adipose fin developmental patterns. (a) Adipose fins (ad.f) develop either *de novo* as a bud following regression of the LMFF or from domain of the LMFF. (b) Phylogenetic distribution of the two developmental patterns. The simplified phylogeny of teleost fishes is based on ^68^ and modified to show recent phylogenetic hypotheses of Otophysan intra-relationships^69, 70^. Black branches indicate lineages that are observed with adipose fins or where the fin is estimated to have occured^3^. Species-level data and associated references are provided in Suppl. Table 1.

### Adipose fin development in Corydoras aeneus

*Corydoras aeneus* (Suppl. Fig. 1) exhibits fold-­associated adipose fin development. Prior to adipose fin development in *C. aeneus*, the LMFF appears undifferentiated between the dorsal and caudal fins (Fig. 2 a). Once the larvae have grown to approximately 8 mm standard length (SL), the LMFF begins reducing both immediately posterior to the dorsal fin and anterior to the caudal fin, and a condensation forms at the future site of the adipose fin, midway along the proximodistal axis of the LMFF (Fig. 2 b). At the anterior boundary of this condensation, an ossification forms that will become the adipose fin spine (Fig. 2 c). The ossification is unpaired, positioned on the midline, and it grows both proximally and distally, parallel to actinotrichia in the LMFF (Fig. 2 d-­f, Fig. 3). Once the spine has extended proximally to just dorsal to the epaxial musculature, its base widens into a saddle shape that wraps laterally around the trunk musculature. Three scutes develop anterior to the adipose fin spine, their order of ossification proceeding from posterior to anterior (Fig. 3 g-­l). The LMFF continues to reduce, leaving a domain posterior to the adipose fin spine that will constitute the membrane of the adipose fin. The LMFF has finished reducing and the adipose fin has fully developed by the time *C. aeneus* reach 1.2 cm SL. Odontodes, small dermal denticles, develop on the scutes of *C. aeneus*^36^. In the adipose fin, odontodes begin mineralizing before the adipose fin spine has ossified (Fig. 3 b).

**Figure 2.**
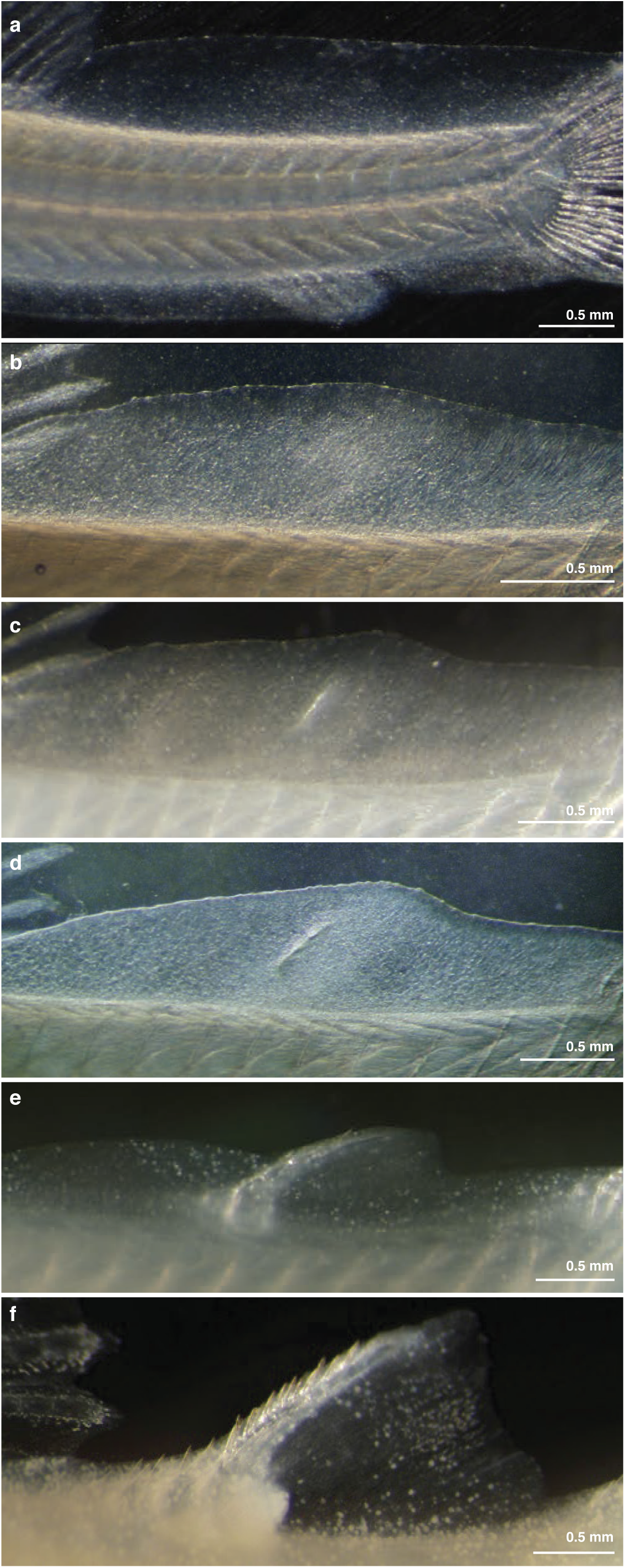
Gross anatomy of adipose fin development in *C. aeneus*. (a) Prior to adipose fin development the LMFF is uniform along its length. (b) The LMFF begins reducing posterior to the dorsal fin and anteri- or to the caudal fin and a condensation is visible where the adipose fin will develop. (c) Ossification is initiated at the anterior portion of the condensation, and the ossification is oriented parallel to the actinori-chia in the LMFF. (d-f) Ossification extends both proximally and distally as the LMFF continues to reduce, and a domain of the LMFF is retained that will become the adipose fin membrane.

**Figure 3.**
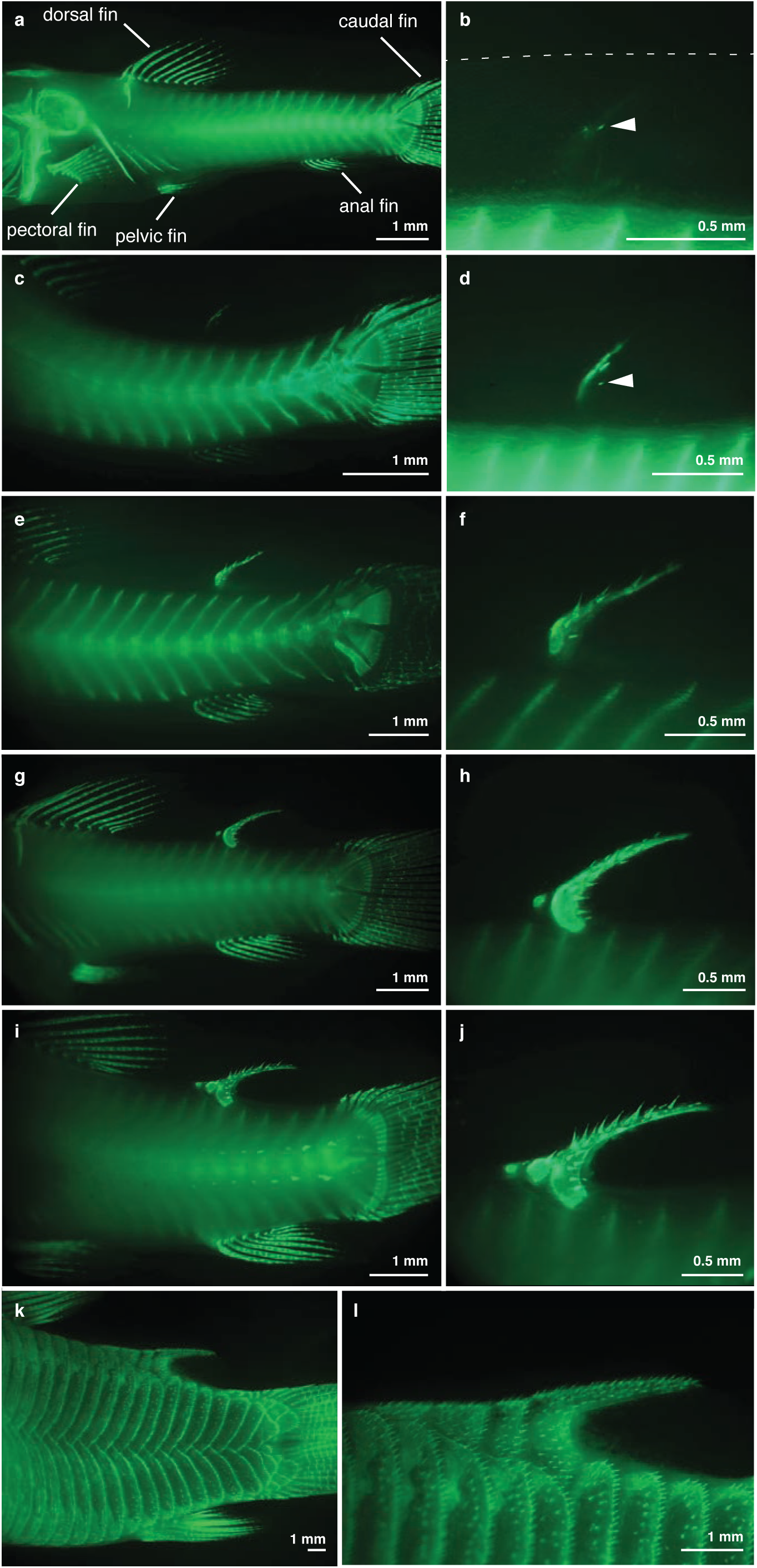
Development of *C. aeneus* adipose fin skeleton. Fixed specimens were stained with calcein to reveal ossification patterns for the adipose fin spine, and odontodes, which are indicated in (b) and (d) by arrow heads.

The adipose fin of *C. aeneus* has extensive sensory innervation^33^. Therefore, we also studied the ontogeny of sensory anatomy and innervation in the adipose fin. Before adipose fin development is observed, the LMFF is innervated by nerve fibers entering from the midline and from the trunk epithelium, and these nerves terminate as free nerve endings (Fig. 4 a). Nerve fibers persist while the LMFF reduces, extending to the LMFF’s distal margin (Figs. 4 b, c and 5 a, b). Adipose fin-­associated nerve fibers are observed posterior to the fin spine by the time the spine has extended to reach the trunk musculature (Fig. 5 a, b). These fibers appear to originate from the dorsal rami of the spinal cord and dorsal projections of the recurrent ramus of the facial nerve, which extends into the caudal fin.

**Figure 4.**
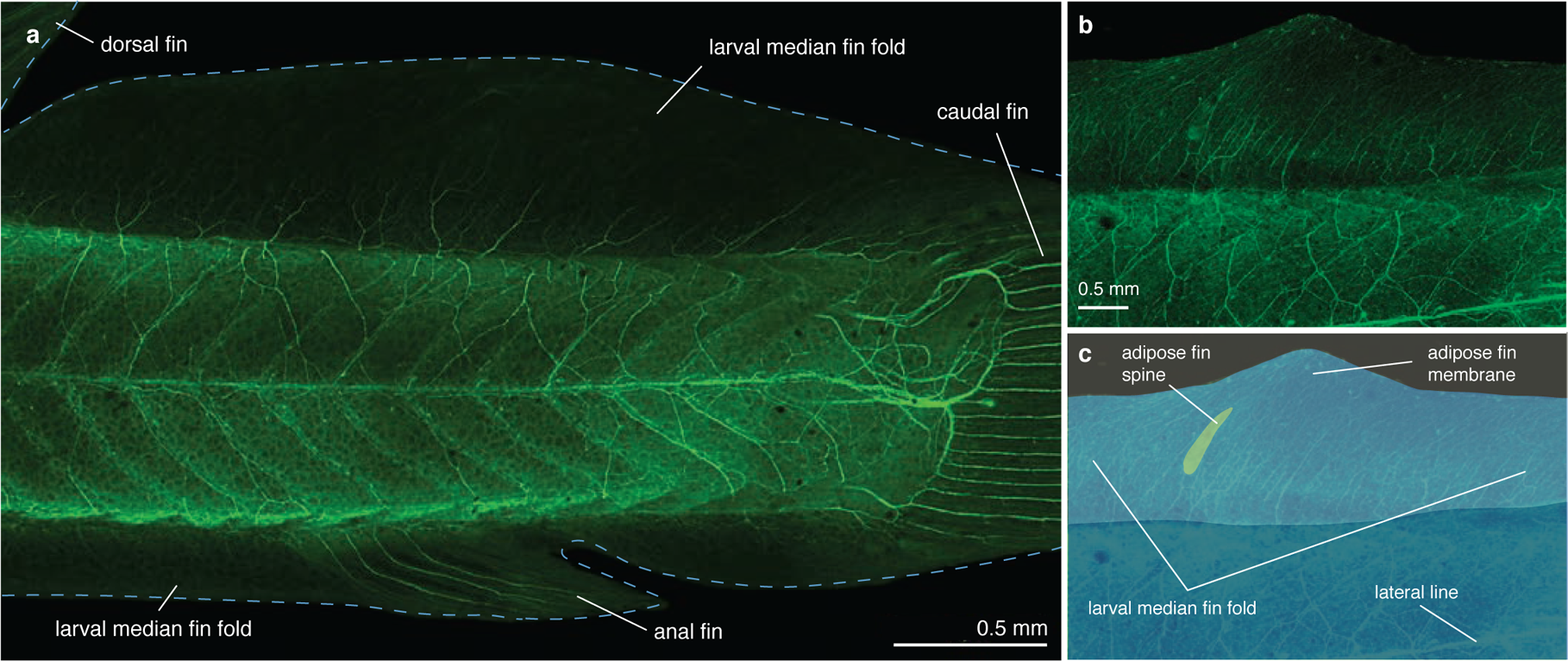
The larval median fin fold contains sensory innervation. Confocal images of larval *C. aeneus* immunostained with anti-acetylated tubulin to label nervous tissue show nerves entering the fin fold and terminating as free nerve endings (a) prior to differentiation of the adipose fin spine. (b,c) Nerves in the LMFF are of a similar organization after the fin spine has begun developing and before the spine has grown to reach the epaxial musculature.

**Figure 5.**
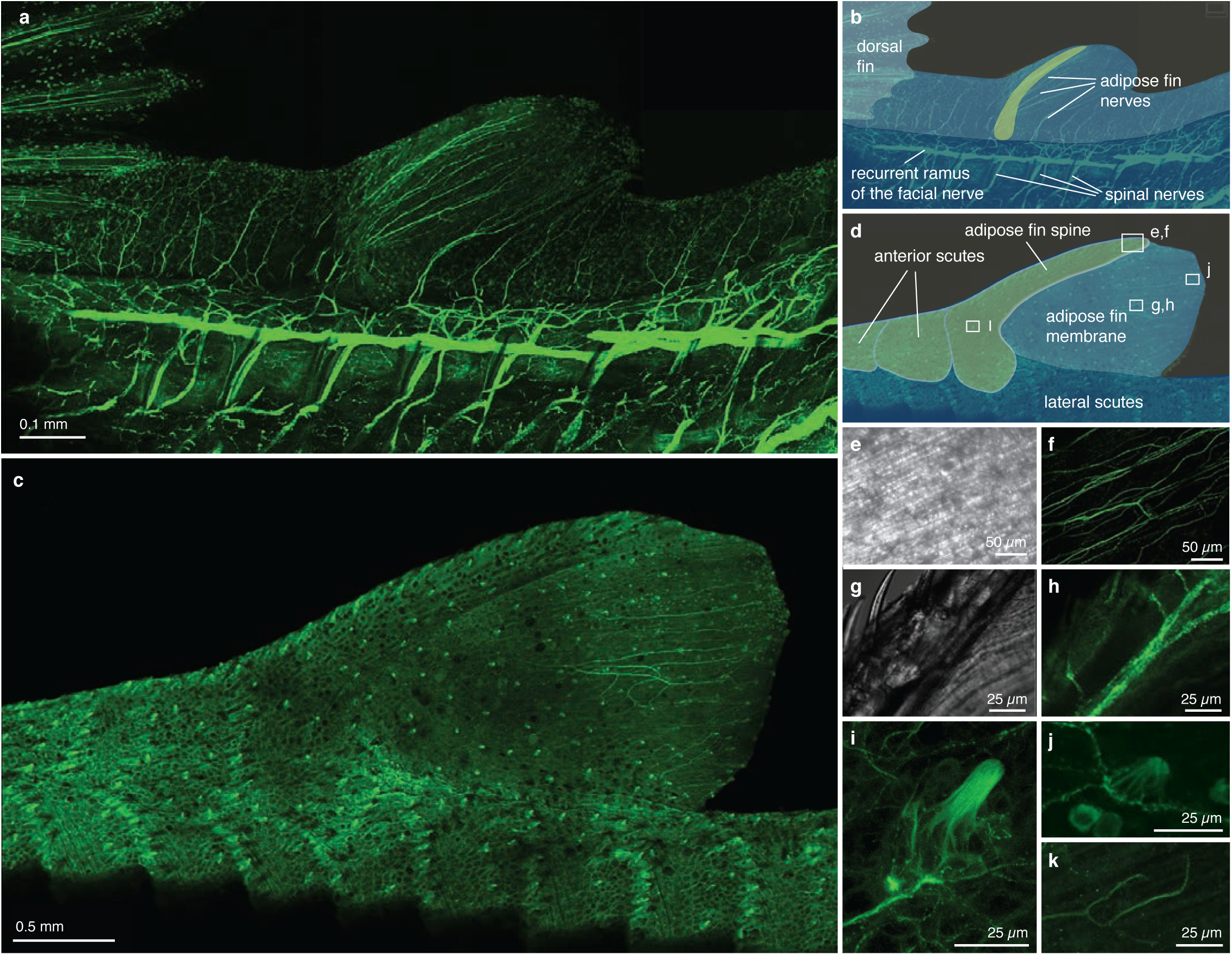
Adipose fin-associated sensory nerves are observed after the fin spine has grown to reach the epaxial musculature. (a,b) Left side of the specimen has been dissected to expose dorsal rami of the spinal cord and the recurrent ramus of the facial nerve. (c,d) Sensory cells and innervation are observed on the fully developed adipose fin. (e, f) Transmitted light and immonofluorescence images show nerves in the fin membrane run parallel to actinotrichia in the fin membrane. (g, h) Transmitted light and immono-fluorescence images showing nerves passing anteriorly through the adipose fin spine and ramifying to extend branches posteriorly into the fin membrane. (i) Superficial neuromasts are observed on the adipose fin spine. (j) Taste buds are observed on the adipose fin membrane. (k) Nerves in the adipose fin membrane also terminate as free nerve endings. Boxes in (d) indicate the approximate position of images in subsequent panels. Panels (i,j) are the same specimen as (c); panels (e-h, k) are of another specimen.

Most of the adipose fin innervation enters the fin as a ramus immediately posterior to the adipose fin spine that ramifies repeatedly to extend branches posteriorly into the adipose fin membrane (Fig. 5 a, b). Additional nerves enter the adipose fin membrane posterior to the fin spine and do not clearly correspond to somatic boundaries. Nerve fibers in the adipose fin membrane are organized approximately parallel to the actinotrichia (Fig. 5 c-­f). Nerve branches also pass anteriorly through the adipose fin spine (Fig. 5 g, h).

Two kinds of putative sensory structures, diagnosed by their morphologies, are observed on the adipose fin of *C. aeneus.* The first kind, observed on the fin spine, are columnar projections with a rounded apex and a base composed of a cluster of globular cells (Fig. 5 i, Suppl. Fig. 2). This morphology is consistent with previous descriptions of superficial neuromasts^37^. These structures are also observed on the lateral scutes of *C. aeneus* and can be distinguished from lateral line sensory cells, which have a filamentous tip (Suppl. Fig. 3). The second kind of sensory structure, observed on the adipose fin membrane, are hillock shaped and lack a pronounced apex (Fig. 5 j). These are diagnosed as taste buds by their size, shape, and distribution^38-­40^. In catfishes, taste buds are distributed across the body and commonly found on both the trunk and fins, including the LMFF and adipose fin. These structures are composed of a cluster of globular cells and are innervated by branches of the adipose fin nerves (Suppl. Fig. 4). Nerves in the adipose fin membrane also terminate as free nerve endings (Fig. 5 k).

## Discussion

Comparative studies of development can inform plesiomorphic conditions and constrain hypotheses of how novel structures originated^41^. In this study, we analyzed the development and diversity of adipose fins to understand their evolutionary origin. These data show that novel appendages can evolve by individuation of a domain of a plesiomorphic fin fold. Additionally, adipose fins that develop from the LMFF can evolve skeleton^3, 32, 42^, musculature^31^, and sensory anatomy (here and ^43, 44^). For example, in *C. aeneus* the domain of the LMFF that contributes to the adipose fin is transformed over ontogeny by the migration and growth of new fin-­associated tissues into the territory. Collectively, these observations show the evolutionary trajectory outlined in the MFF hypothesis—partitioning of an existing median fin fold and transformation of such a fin into a structurally and functionally complex appendage—is a viable route by which novel fins can originate.

Adipose fins evolved independently in Euteleostei and Otophysii^3^. Euteleosts uniformly exhibit fold-­associated adipose fin development, indicating that adipose fins originated in this clade, at least in part, by the retention of a larval character that would have primitively been reduced over ontogeny. For example, in *Polyodon spathula* (Acipenseridae) the LMFF reduces fully between the dorsal and caudal fins in development^23^. In otophysans, adipose fins develop in two ways: fold-­associated development in Siluriformes, and *de novo* outgrowth in Characoidei. These two groups are sister to one another, and data is unavailable for Citharinoidei, the clade sister to that total group that also have adipose fins. Therefore, we cannot currently resolve the primitive pattern of adipose fin development in otophysans. The occurrence of both developmental patterns in Otophysii reveals evolutionary lability between these two states, although the polarity of change is unresolved, and this raises questions of how adipose fins are positioned along the anteroposterior axis in ontogeny.

Fin positioning depends upon several processes: genes expressed in the mesoderm (either lateral plate^45^ or paraxial^46, 47^) establish the fin territory, cells in this territory can migrate and contract asymmetrically to form a mysenchymal bud^48^, and differential growth of the trunk and fin can affect its eventual adult position^26, 49, 50^. Some adipose fins appear to be positioned by an additional mechanism: differential patterns of apoptosis within the LMFF. How this is regulated, or how it relates to positioning in adipose fins that develop *de novo* as buds is unclear. Future work should characterize adipose fin development in Citharinoidei, to understand whether *de novo* or fold-­associated development is primitive in otophysans. Additionally, characterizing the developmental genetic mechanisms that underlie positioning in Characoidei and Siluriformes adipose fins would clarify how lineages transition between these two apparently disparate developmental patterns. Developing such a model would allow for detailed hypotheses of how transitions predicted by the MFF hypothesis occurred in early chordates.

Adipose fins have evolved anterior dermal spines from midline scutes three times independently (*i.e.*, *Sisor spp*. (Sisoridae), Amphilidae, and Lorocaricoidea, which includes *C. aeneus*)^3^. Scutes can be distinguished from other postcranial dermal plates by the presence of hyaline, a superficial hypermineralized tissue^51, 52^. In *C. aeneus*, scutes are positioned laterally, on the flank, as well as on the dorsal midline anterior to the adipose fin. Lateral scutes begin developing at the mid-­region of a population of mesenchymal cells and not adjacent to an epithelial-­mesenchymal boundary^52^. Similarly, the fin spine of *C. aeneus* appears to ossify at the core of the LMFF and medial to the actinotrichia and not adjacent to the epithielium. By contrast, lepidotrichia develop from mesenchyme in the space between the basal lamina of the epithelium and actinotrichia;; they are, therefore, comprised of paired elements, the hemitrichia^18^. The medial, unpaired position of the adipose fin spine has consequences for the position of sensory nerves, discussed below.

Although both the adipose fin spine and lepidotrichia develop by the migration of osteogenic cells along actinotrichia—bone differentiation and growth proceeding parallel to the orientation of actinotricha^53^—these two skeletal types differ in the site of initial ossification and direction of skeletal growth. The adipose fin spine of *C. aeneus* begins ossifying midway along the proximodistal length of the LMFF and extends both proximally and distally. This is in contrast to the general pattern of ossification in vertebrate appendages, where skeleton usually differentiates and grows in a proximal-­to-­distal direction^54^. For example, with a single exception (*i.e.*, a pimelodid catfish that has evolved a rayed adipose fin^32^), lepidotrichia development is initiated proximally and extends distally^17, 55^.

Previous descriptions of odontode development in *Corydoras* observed their differentiation only after the mineralization of associated scutes and fin rays to which they attach^36^. It was, therefore, suggested that odontode ossification depends upon prior ossification of its attachment site^36^. However, our data show that ossification of odontodes can precede scute ossification (Fig. 3 b), indicating the presence of a well-­developed stratum compactum in the fin spine mesenchyme at these stages^51^.

The organization of innervation in the adipose fin of *C. aeneus* is distinct from what has been described in other actinopterygian fins. In rayed fins, sensory nerves enter the fin medially between the paired hemitrichia, sending branches into the fin membrane through the joints between fin ray segments^56, 57^. By contrast, in the adipose fin of *C. aeneus* a large nerve branch enters the fin posterior to the adipose fin spine. This nerve bundle maintains an association with the spine, traveling along the length and sending branches both anteriorly through canals in the spine, and also posteriorly into the fin membrane. The posteriorly directed nerve branches run approximately parallel to the actinotrichia in the fin membrane. Additional nerves enter the fin posterior to the adipose fin spine but do not follow a clearly segmented pattern, as is observed in fins with a series of lepidotrichia (*e.g.*, dorsal, anal, caudal fins, Fig 4 and 5). Future work should identify the cues that guide axonal growth into this territory and compare them to the mechanisms by which sensory anatomy is reorganized in other fins.

For example, the paired pectoral fins of *D. rerio* first develop a non-­skeletonized fin fold that contains sensory innervation. These nerves are not organized in a strictly proximodistal orientation, but instead form a reticulate network that is reshaped over development;; in the adult morphology, sensory nerves are associated with lepidotrichia^57^. Similarly, the LMFF, which is a passive appendage (*i.e.*, lacking musculature control), contains sensory nerves. These nerves are likely from Rohon Beard cells, primary sensory neurons with peripheral axons that extend along the trunk epithelium, into the caudal fin fold, and likely also extend into the dorsal LMFF^58^. These cells undergo apoptosis over ontogeny and do not retract during degradation^59^. In the caudal fin, these are eventually replaced by the dorsal root ganglia.

The adipose fin of *C. aeneus* is innervated by both the recurrent ramus of the facial nerve and by dorsal rami of the spinal cord. We observe both sources projecting nerves dorsally into the adipose fin domain, and other fins have been described with sensory innervation from both cranial and spinal sources^60, 61^. In catfishes, taste buds are distributed across the body and commonly found on both the trunk and fins, including the LMFF and adipose fin^38-­40^. Taste buds on the trunk have been exclusively described as innervated by the recurrent ramus of the facial nerve^40^. Superficial neuromasts have been described previously on the caudal fins of various fishes^37, 62, 63^, but not on adipose fins. Although we were able to observe nerves entering the adipose fin and extending anteriorly through the fin spine, we were unable to determine whether they terminated upon the superficial neuromasts.

The adipose fin of *C. aeneus* is proprioceptive, able to detect the movement and position of the fin membrane^33^. Mechanosensation might be achieved through by either the taste buds or free nerve endings in the fin membrane. In the channel catfish, *Ict*alurus *punctatus*, the nerves that terminate on extra-­oral taste buds on the flank are mechanosensitive^64^;; however, it is unknown how this is achieved. Extra-­oral gustatory cells might detect both mechanical and chemical signals, or the nerves that innervate these cells might also terminate on yet unidentified mechanosensory endings.

The adipose fin of the brown trout, *Salmo trutta* (Salmonidae), a euteleost, is also invested with sensory nerve fibers and also hypothesized to detect fin movement^43, 44, 65, 66^. These nerves terminate upon associated astrocyte-­like cells, which are hypothesized to detect the deformation of collagen fibers that span the left-­and-­right sides of the adipose fin^43, 44^. The organization of collagen is similar to what has been described in the LMFF of *D. rerio*^67^, indicating that many structural components of the LMFF are retained over ontogeny and that sensory anatomy is reorganized over ontogeny in this lineage, similar to *C. aeneus*. Sensory innervation might, therefore, be among the first features to change when a new median fin originates by partitioning of a LMFF.

The earliest vertebrate fins lack evidence for muscular attachment^5, 6^ and were likely passive appendages. Typically, discussions of how these fins originated focus on hypotheses of selection for the functions of stabilization, control, and thrust generation^11^. Given that passive fins (*e.g.*, LMFF and adipose fins) are invested with sensory anatomy^6, 41^ and can be proprioceptive^33^, we argue that selection for flow detection or other sensory functions could also have shaped the evolution of fins in early vertebrates.

### The origins of median fins

The median fin fold hypothesis is the leading model for how differentiated fins originated in vertebrates. However, there is limited evidence from the paleontological record of a transformational series between a continuous MFF to spatially separated fins by reduction of the MFF in certain positions. Claims that adult fins develop from the LMFF—which, if the MFF is understood as homologous to the LMFF, could provide recapitulist evidence for the MFF hypothesis—are similarly weak (*e.g*., numerous *D. rerio* mutants with malformed LMFFs develop normal adult fins^25^, and most structures that comprise the adult fin are not derived from LMFF tissues^20, 26, 27^).

The data presented above on the development of adipose fins provide indirect evidence for the MFF hypothesis. Specifically, they demonstrate that a median fin fold, structurally rudimentary and undifferentiated along its anteroposterior length, can be partitioned into an individuated fin. Additionally, they show that fins that develop primitively from LMFF can be transformed by association of new tissues into this territory such that a nascent fin can evolve to become structurally and functionally complex. Indeed, some catfishes have redeployed the developmental program for lepidotrichia into their adipose fins, such that the adipose fin superficially looks like a duplicated first dorsal fin (*e.g.*, *Mochokus* (Siluriformes))^3^. Although indirect, adipose fin development and diversity might provide the clearest evidence that the MFF hypothesis describes a viable route by which novel fins can originate.

## Methods

Albino *C. aeneus* (n=18) were donated for breeding by NBM Aquatics (Chicago, IL) and housed at The University of Chicago in 10 g tanks at 20°C with a standard seasonal light/dark cycle. Fish were conditioned for breeding on a diet of live blackworms, *Lumbriculus variegatus*, and stimulated to breed with 50% water changes. Females deposited eggs on the aquarium glass, and embryos were transferred by hand to 1 L containers of aerated tank water treated with three drops of the antifungal Methylene blue (VWR International, West Chester, PA, USA;; 0.5% diluted in Hank’s solution). Water was changed 30% daily until three days post-­hatching, at which point larvae were transferred to mesh boxes suspended in the adult tanks and raised on a diet of dried shrimp pellets (OmegaSea, LCC, Painesville, OH).

To generate a developmental series, larvae were collected through one month post-­ hatching. Specimens acquired through the pet trade were used for adult stages. Specimens ranged in size from 0.5–2.7 cm SL. Animals were euthanized with MS222 (Tricaine methanesulfonate, Sigma-­Aldrich, St. Louis, MO) at a concentration of 0.5 g/L, fixed in 4% paraformaldehyde at 4°C for 3 days on a rocker, and then stored in 100% methanol at 4°C. Animal care and euthanasia protocols were approved by the University of Chicago’s Institutional Animal Care and Use Committee.

Calcein, which fluoresces when bound to calcium, was used to characterize skeletal development (Sigma #: C0875, Sigma Chemical Co., St Louis, MO). Fixed specimens (n=6) were immersed in an aqueous solution of 50% methanol and 1% Calcein for 36 hours at 4°C on a rocker and then rinsed three times in 50% methanol prior to imaging. Photographs of the developmental series and of calcein stained specimens were collected using an Olympus DP72 camera mounted to a Leica MZ10 microscope (Leica Microsystems, Wetzlar, Germany) using the software CELLSENS ENTRY v.1.2 (Build 7533) software (Olympus Corporation, Tokyo, Japan).

To characterize neuroanatomy, fixed specimens (n=28) were antibody stained using methods adopted from Thorsen and Hale [1]. Nerves were labeled using the primary antibody Mouse monoclonal anti-­acetylated tubulin (Sigma-­Aldrich), and the secondary antibody goat anti-­ mouse antibody conjugated with fluorescein (Jackson ImmunoResearch Laboratories, West Grove, PA). Specimens were stained both undissected and following the removal of skin and myomeres from the left side of the body to expose nerves entering the fin. Antibody stained specimens were imaged with a Zeiss LSM710 confocal microscope (Carl Zeiss Inc., Thornwood, NY, USA). Using the LSM710 software, images with extended depth of field were generated using the ‘maximum intensity projection’ function on a z-­stack of images. Also using the LSM710 software, large samples were imaged by ‘tiling,’ wherein adjacent z-­stacks with 10% overlap were combined into a single image. Brightness and contrast have been uniformly adjusted for images presented in Figures 4 and 5, so that they are consistent across samples and microscope settings.

## Acknowledgements

Thank you to M Schauer at NBM Aquatics for guidance on the acquisition, husbandry, and breeding of *Corydoras*. Thank you to MI Coates, WL Smith, BR Aiello, F Stabile and D Krentzel for feedback on the manuscript. This research was supported by the following: National Science Foundation, IGERT grant no. DGE-­0903637 and GRFP to TAS;; the Office of Naval Research grant N00014–0910352 to MEH;; National Institute of Health grant HD072598 to RKH, the American Society for Ichthyologists and Herpetologists, Raney Award;; The University of Chicago, Hinds Fund.

## Author contributions

T.A.S. conceived of the project, collected and analyzed data, and wrote the paper. R.K.H. and M.E.H. contributed to the design of the study, discussed and interpreted results, and helped to write the paper.

## Data availability statement

The datasets generated during the current study are available from the corresponding author on reasonable request.

---

**Suppl. Table 1:**
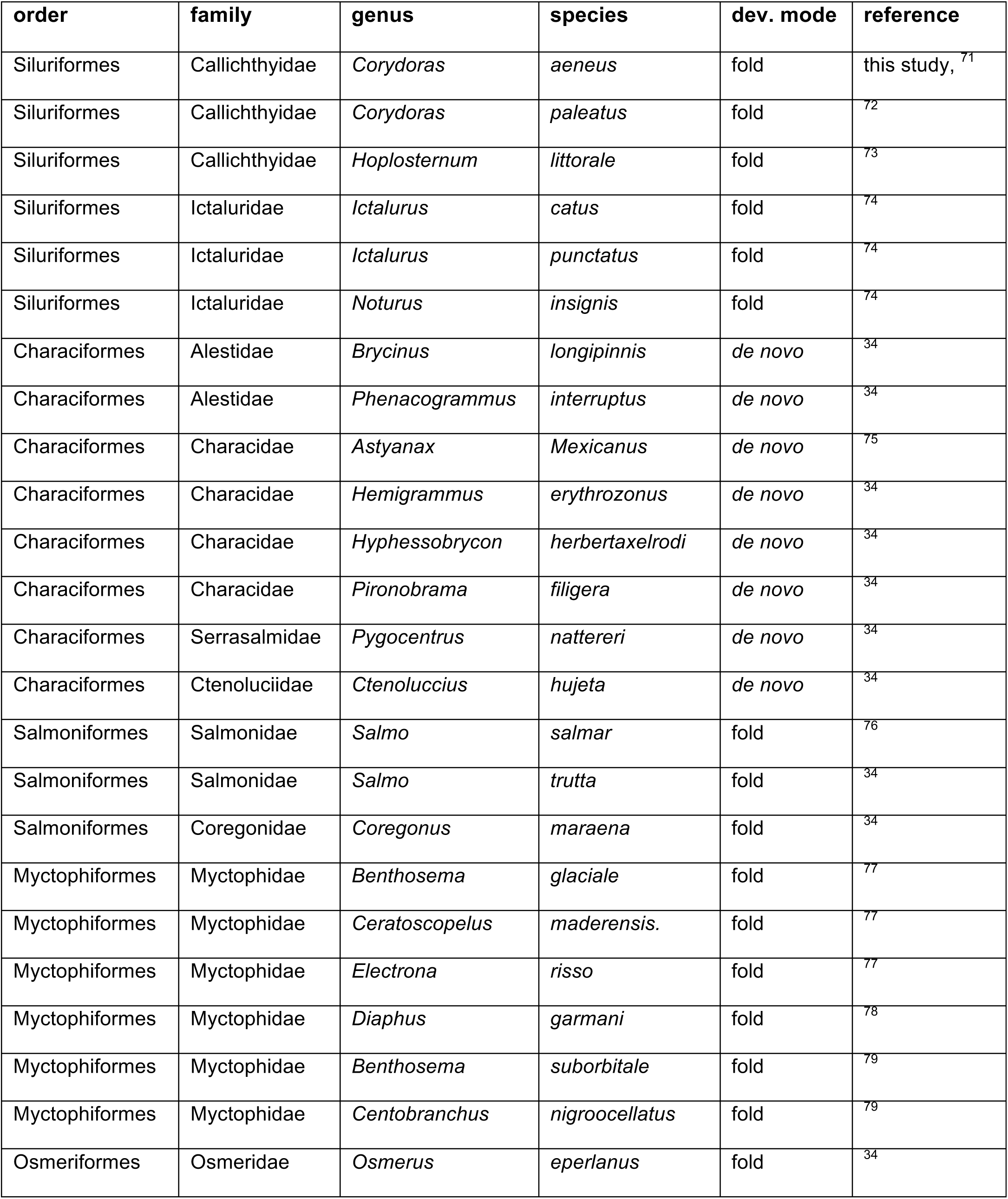
Data aggregated from the literature on adipose fin developmental diversity. Information on adipose fin development was found in one study of the fin’s development 34, the taxonomic keys of larval fishes, and in developmental staging papers. The information is summarized in Fig. 1.

**Supplementary Fig. 1.**
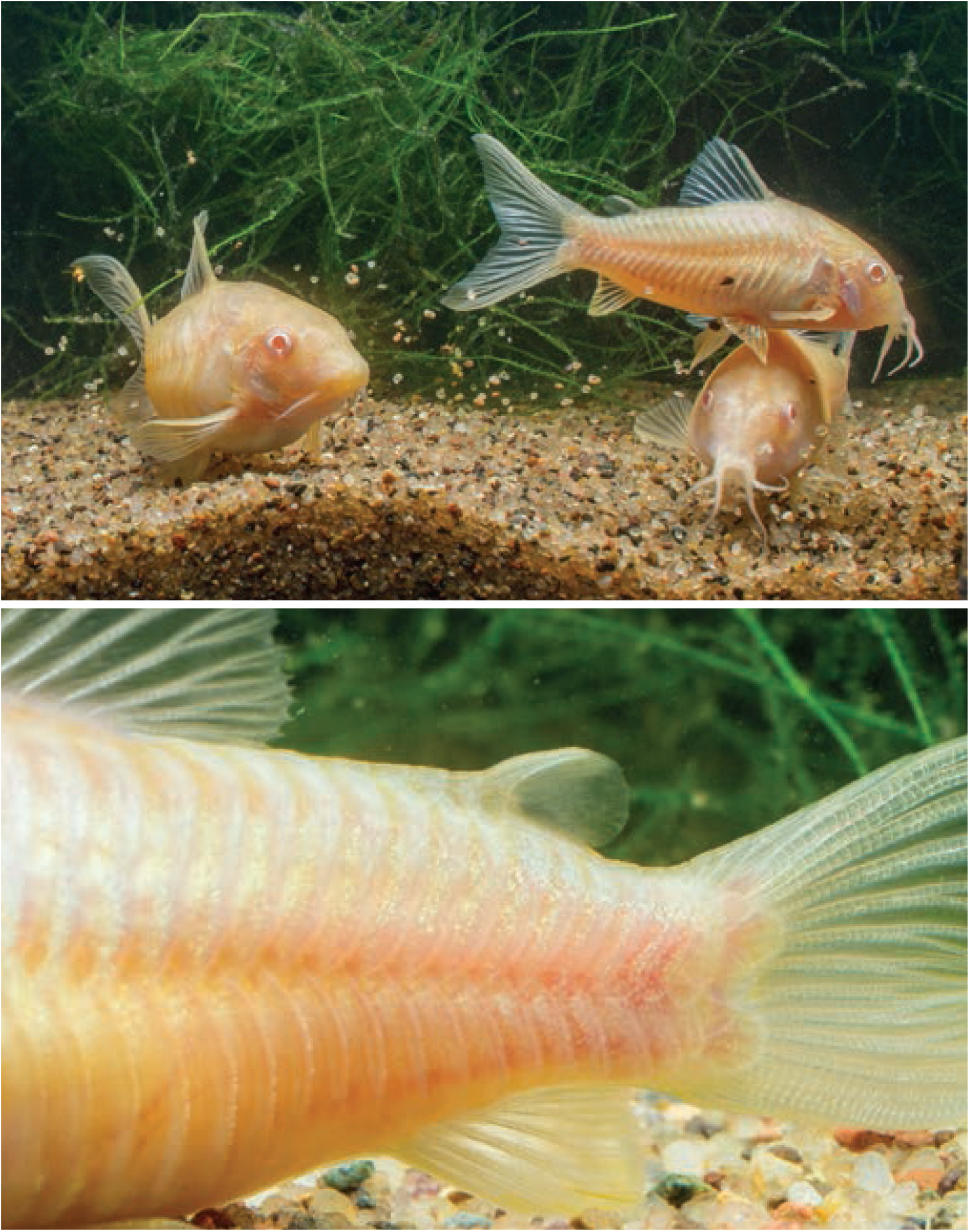
Photographs of adult *C. aeneus* by Yen-Chyi Liu.

**Supplementary Fig. 2.**
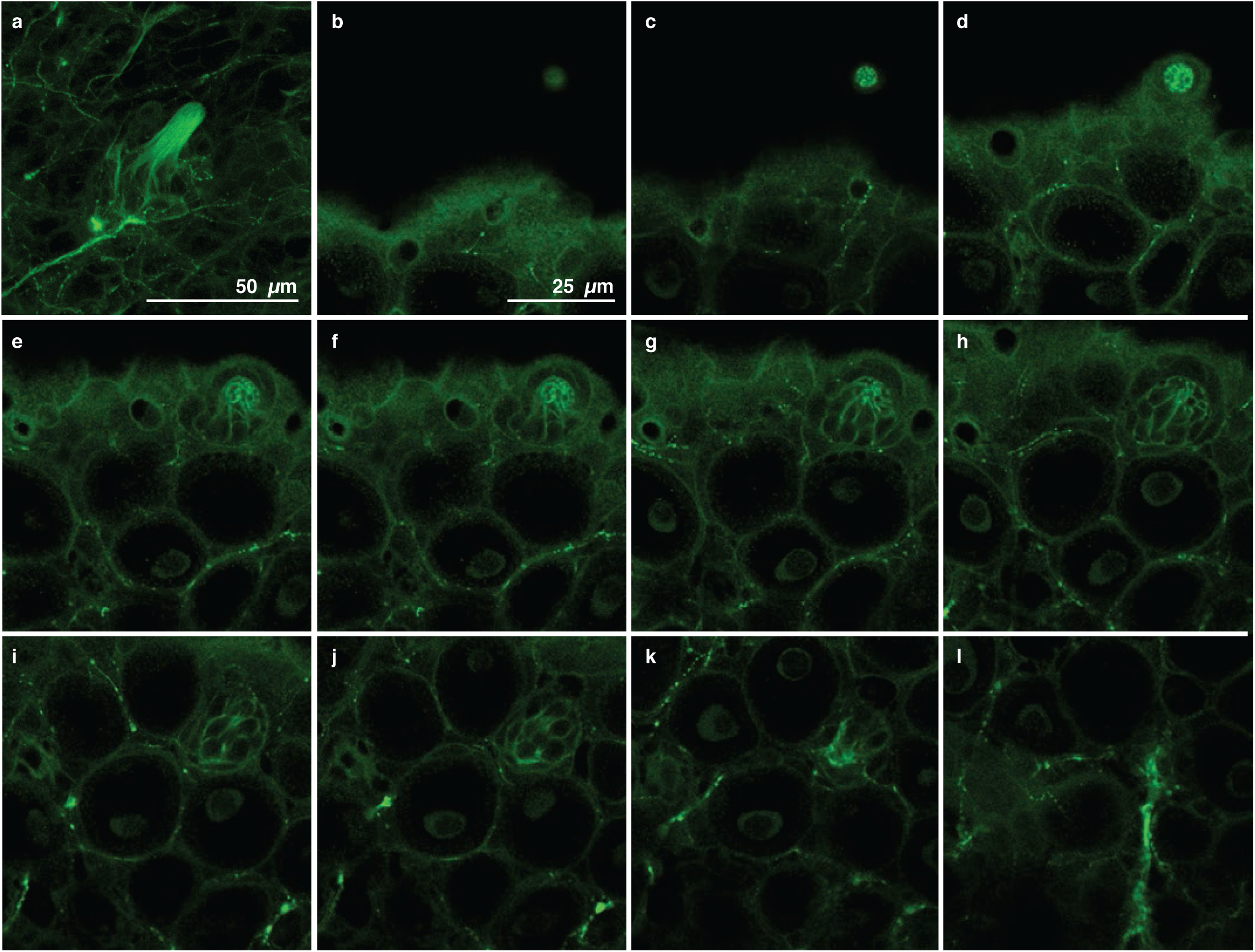
Superficial neuromasts on the adipose fin spine and lateral scutes of *C. aeneus*. (a) Maximum-intensity projection of a superficial neuromast on the adipose fin spine. Panels (b-l) are of a neuromast on a lateral scute, which is oriented perpendicular to the image plane. Panels are from apex (b) to base (l). Depth between panels is 2.7 μm.

**Supplementary Fig. 3.**
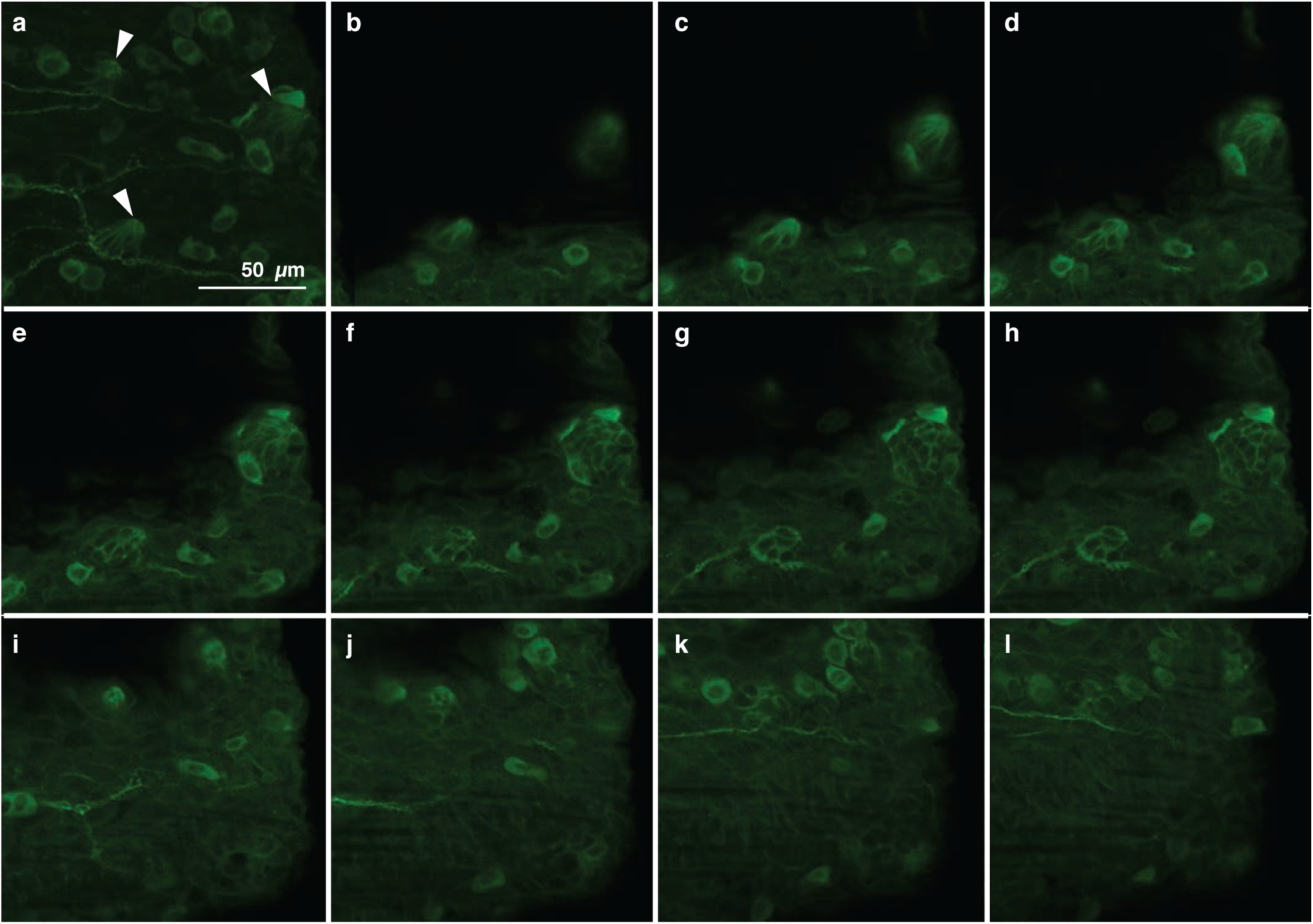
Anatomy of a taste buds on the adipose fin of *C. aeneus.* (a) Maximum intensity projection showing three taste buds, indicated by arrowheads, and associated nerves. The taste buds are oriented perpendicular to the image plane, and panels are from apex (b) to base (l). Adipose fin afferents, which run parallel to actinotrichia in the membrane, terminate upon taste buds. Depth between panels is 3 μm.

**Supplementary Fig. 4.**
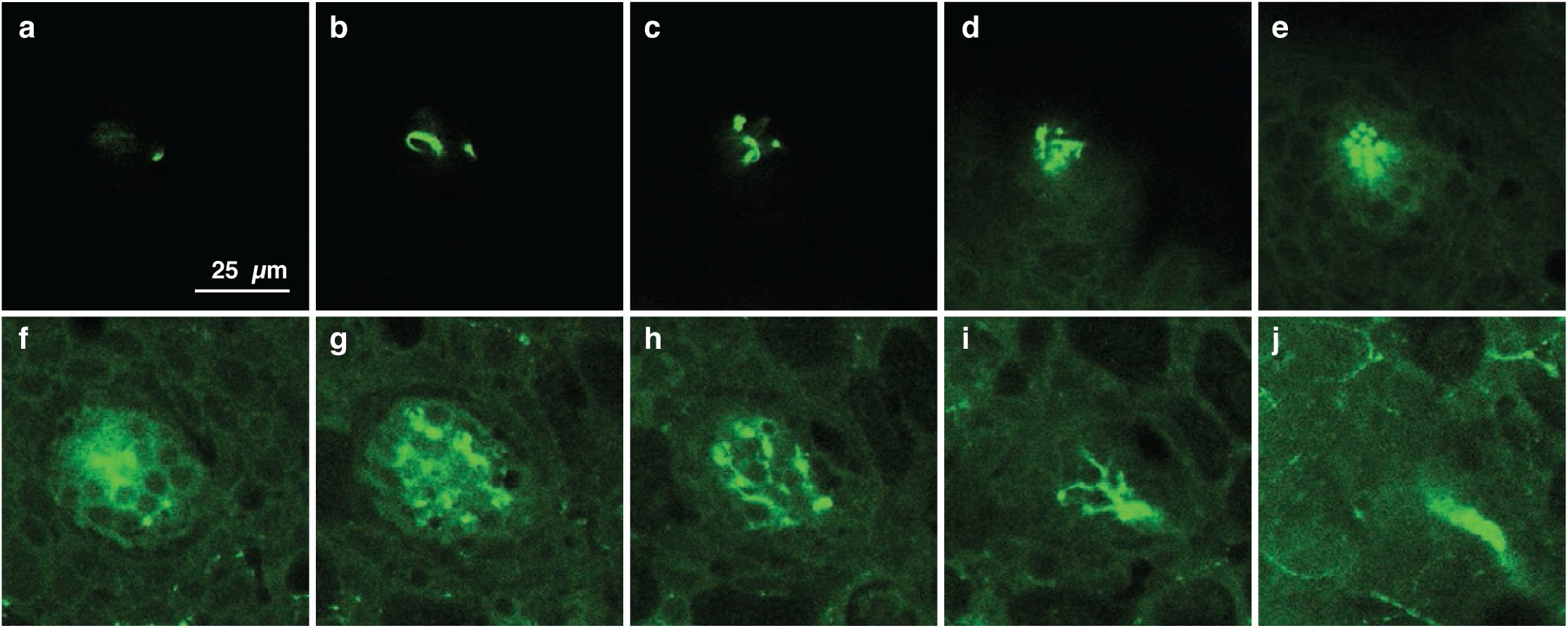
Anatomy of a lateral line cell of *C. aeneus.* Panels (a-j) are of a lateral line cell positioned ventral to the adipose fin on the left side of the body; anterior is left. The lateral line cell is oriented perpendicular to the image plane, and panels are organized from apex (a) to base (j). The apex of the lateral line cell is characterized by a filamentous tip. Depth between panels is 5.5 μm.

